# Sleep in People Experiencing Homelessness Under Different Conditions and Seasons

**DOI:** 10.1101/2025.01.23.634551

**Authors:** Alicia Rice, Leandro P. Casiraghi, Cristina Gildee, Zack W. Almquist, Amy Hagopian, Melanie A. Martin, Horacio O. de la Iglesia

## Abstract

Poor sleep represents a central health disparity for people experiencing homelessness, and any intervention to alleviate the impacts of homelessness should aim to improve sleep. We measured actimetry-based sleep in homeless adults spending their nights in four types of shelters in Seattle, WA, during the summer and winter. Homeless participants experienced more sleepless nights than housed participants in both seasons. During the summer sleeping nights, homeless participants experienced sleep patterns similar to housed subjects, but during the winter, their sleep duration was up to 1.5 hours shorter. Similarly, sleep quality, determined through sleep variability index, activity during the night, and intraindividual variability of sleep parameters, was poorer during the winter in homeless than in housed participants. Our study demonstrates the feasibility of using objectively measured sleep as a proxy for assessing the value of specific interventions to improve living conditions in people experiencing homelessness. Given the bidirectional relationship between sleep and both physical and mental health, our study reveals the health inequities of chronic sleep disparity for those living outdoors.

**Significance statement:** Sleep disparities are central to the poor physical and mental health experienced by houseless people. Here we show that measured sleep through wrist actigraphy captures different aspects of this sleep disparity and represents a useful, objective metric of the impact of interventions to improve living conditions among people experiencing homelessness.

## Introduction

The crisis around the number of people experiencing homelessness has reached alarming numbers throughout the world, moving the United Nations (UN) General Assembly to adopt resolution 76/133 recognizing homelessness as a major human rights issue in 2021.

Homelessness riddles all nations, including highly developed economies like the United States (US). One of the most hard-hit areas in the US, King County, WA (greater Seattle area), is the 12^th^ most populated county (2.267 million people), which has recorded the third highest homeless count, driven by massive increases in the cost of living. The primary US method for counting people experiencing homelessness on a national scale is based on the so-called Point-in-Time (PIT) count, an estimate of people experiencing homelessness conducted by hundreds of local jurisdictions. The 2024 PIT count reported 16,385 people experiencing homelessness in King County, a 23% increase from 2022^1^.

The high incidence of homelessness reveals a crisis in human rights, social justice, and public health, which local governments and non-profit organizations try to alleviate through interventions to reduce the harm to individual health and the local economy. In Seattle, these interventions include multiple forms of “emergency shelter,” including semi-private continuous occupancy shelters, overnight shelters, tiny houses, and sanctioned tent encampments (aka tent cities; see Methods for description of sites). In 2024, it was estimated that 6,586 people were using emergency shelters in King County, with an estimated 100 people sleeping in tiny houses and 3,136 in tents^2^.

Many fundamental building blocks of health are missing or crumbling for people experiencing homelessness, including nutrition, hygiene, security, and comfort. Perhaps the biggest challenge unhoused people face is the lack of a proper sleep environment. Indeed, several survey-based studies have shown that unsheltered people report poor sleep quality and quantity, that this disturbed sleep is associated with negative health outcomes, and that specific factors may causally mediate the link between poor sleep and poor physical and mental health^3–10^. Insufficient and low-quality sleep has a bidirectional relationship with physical and mental disease: poor sleep worsens core symptoms of virtually every physical and mental disorder, and these symptoms decrease sleep quality. Thus, people experiencing homelessness are caught in a vicious circle that is extremely hard to break.

A critical goal of any intervention to cope with homelessness should be to improve sleep, which can serve as a metric of the success of the intervention and also improve health outcomes.

However, typical methods of tracking sleep quality and quantity among people living rough^3,5,7^ have two important limitations: 1) they are based on a self-assessment of sleep quality and quantity, and 2) this assessment takes place as a single questionnaire that asks for a retrospective analysis of sleep. Although these methods are sensitive enough to detect coarse sleep deficiencies associated with homelessness, they lack the resolution to detect smaller differences in sleep timing and quality that may be present between people experiencing homelessness under varying circumstances, such as relatively tiny houses vs. congregate sleeping arrangements in overnight shelters.

Our study sought to measure sleep among people in various sheltered situations, using an objective and consistent measure of sleep: wrist actimetry. Actimetry represents the gold standard for the study of sleep under field conditions because it is the most objective and quantitative measure of both sleep timing and quality under conditions where polysomnography is not feasible. We report the results from the unhoused population in four types of emergency shelter settings in Seattle: a continuous-occupancy shelter, an overnight youth (18-25 years of age) shelter, tiny houses, and a tent city. Because of the large seasonal variation on the influence of the elements at the Seattle latitude, we also compared sleep parameters in the same communities between the summer and the winter. Lastly, we compare seasonal sleep quality among unhoused participants with that of housed participants from a separate mixed-age study sample.

## Results

### Sleep Timing Between Unhoused and Housed Communities During Summer and Winter

We compared four critical sleep timing parameters —onset, offset, duration and midsleep— generated by individuals in each homeless community to sleep parameters in housed individuals during the winter and summer, adjusted by gender. We built a linear model for each season, where coefficient estimates for each unhoused community were compared to the housed individual values (Table S1). There was an effect of gender on sleep onset clock time (females: 22:43, males: 23:54, non-binary: 23:36, trans: 23:21; p = 0.0455) and sleep duration (females: 8hr5min, males: 7hr18min, non-binary: 8hr11min, trans: 6hr18min; p = 0.0158). Fig. 1 shows the sleep timing of unhoused and housed communities during the summer and winter. During the summer (Table S1.a), sleep onset, offset, and midsleep times did not show a significant difference between housed and unhoused communities. Summer sleep duration in tiny house participants (7h1min ± 26min) was 45 min shorter than in housed individuals (7h46min ± 24min; p = 0.0428). Importantly, this duration average does not include sleepless nights; including them would bring the average tiny house sleep duration to 6h35min. Bartlett’s tests indicated that the sleep midsleep variance differed between communities (p = 0.0454), with the overnight shelter showing the lowest variability in midsleep.

**Figure 1.**
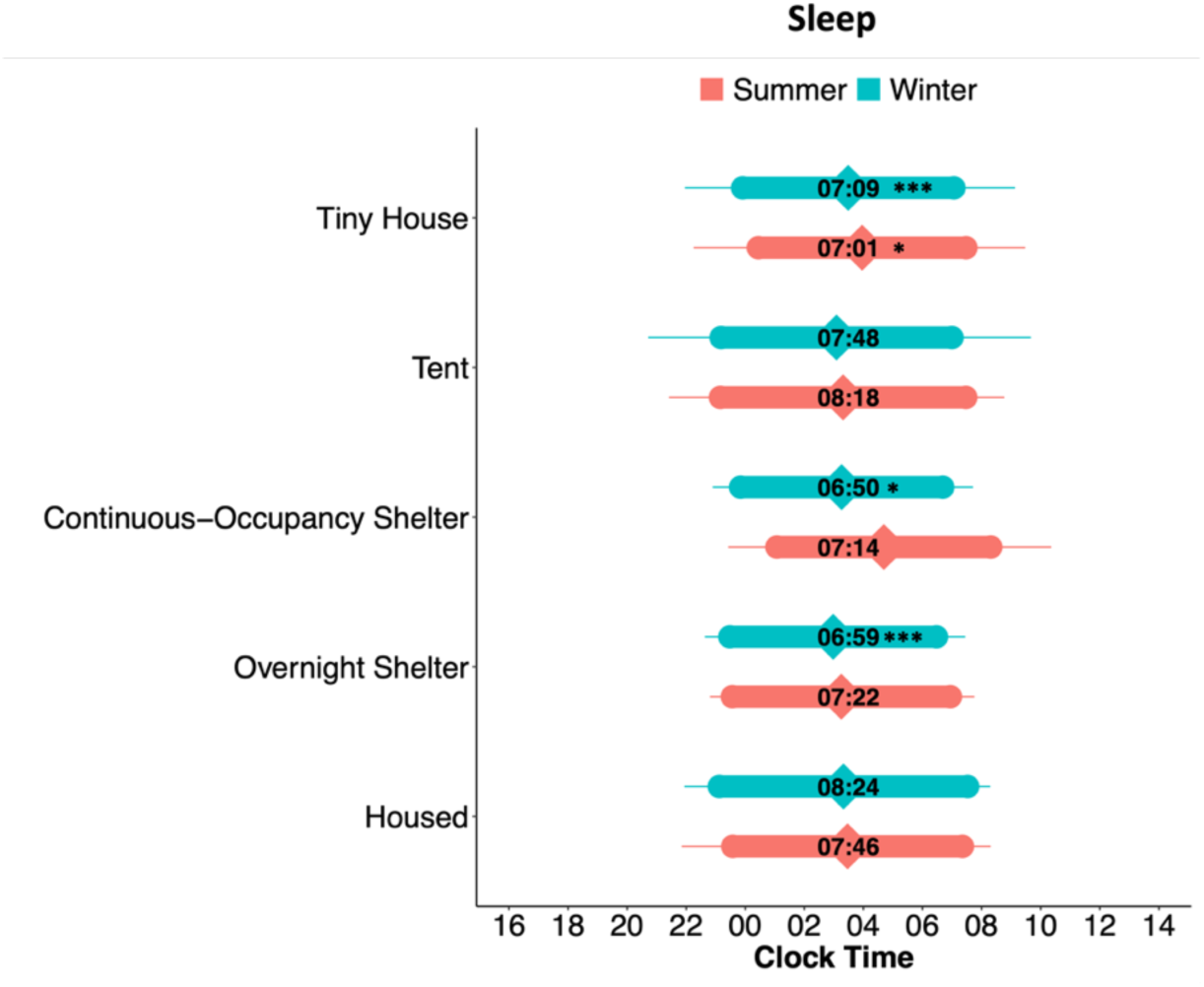
Sleep timing in housed and unhoused participants. Bar plots of average sleep onset, offset, and sleep duration. The diamond shapes within the coloured bars represent the 24-hour clock time of the midpoint of sleep. The numbers within each bar represent the average duration of sleep (e.g., 7:09 in HH:MM for Tiny House). Sleep plots for each community are split between seasons (summer or winter). Error bars represent standard deviations of the mean; note the large standard deviation in tent sleepers during the winter). Symbols represent linear model coefficient values comparing each type of unhoused community to the housed group (seasons compared separately): *p < 0.05; **p < 0.01, ***p < 0.001, difference in sleep duration. See Table S1 for linear model coefficient results.

During the winter (Table S1.b), sleep onset, offset, and midsleep times did not show a significant difference between unhoused sleeping communities and the housed participants. Sleep duration, however, differed during the winter between housed and several unhoused community types. On average, housed participants slept more than an hour and a half longer than participants in the continuous-occupancy shelters (1h34min, 8h24min ± 16min, vs. 6h50min ± 23min, p = 0.0168), 1h25min longer than overnight shelter users (6h59min ± 15min, p = 0.0003), and 1h15min longer than participants in tiny houses (7h09min ± 15min, p = 0.0003). Bartlett’s tests indicated that the time of sleep onset (p = 0.0365), sleep offset (p =0.0021), and midsleep (p = 0.0035) variances were different between all communities, with the lowest variability in the overnight shelter (where entry and exit times are rigid) and housed individuals.

### 24-hour Activity Patterns

We then inspected the average locomotor activity profiles of each community throughout the 24-hour day (activity “waveforms”; Fig. 2). A two-way ANOVA revealed an effect of time of day (p < 0.0001) and an interaction between community type and time of day (p < 0.0001) during the summer. We then compared the morning and evening activity levels within time windows where visual inspection of the means ± standard error of the mean (SEM) suggested differences between communities. This comparison revealed differences in morning activity between communities (Table S2). A comparison of activity during the winter yielded an effect of time (p < 0.0001) and an interaction of community and time of day (p < 0.0001). Differences in activity levels were found during the morning time window (Table S2). Visual inspection of the waveforms suggests that the high activity levels of participants in the overnight shelter drive differences in the mornings and evenings. This high morning and evening activity is preceded and followed by rest, respectively, which is most likely a consequence of the strict schedule of the overnight shelter (doors open at 21:00 in the evenings, and morning departure is enforced before 8:00).

**Figure 2.**
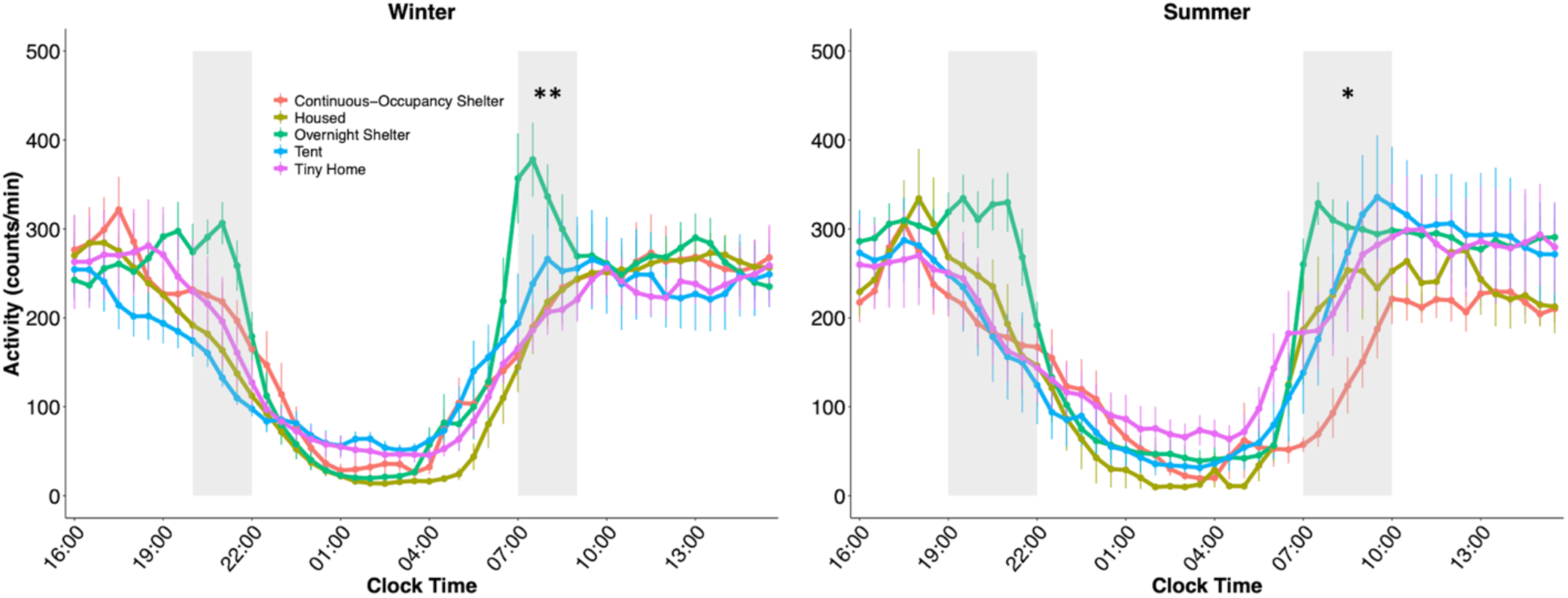
Locomotor activity waveforms in housed and unhoused participants. Points represent participants’ average (±SEM) activity levels for each time point, split between seasons. Grey-shaded areas represent morning and evening time windows during which the summation of activity was compared between communities. *p < 0.05, **p < 0.01, one-way ANOVA analysis.

### Sleep Quality

We examined the number of nights in which no sleep was detected either by wrist actimetry or sleep diary entries (see Fig. S1 and Methods for details). A chi-squared analysis showed a significant difference between communities (p < 0.0001). Out of all recorded nights, the highest number of sleepless nights was found in tent (n = 26 nights) and tiny house participants (n = 37 nights), and they occurred both during the winter and summer.

We then used three different estimates of sleep quality that can be derived from wrist actimetry. First, we measured the sleep regularity index (SRI), an estimate of how regular sleep is that calculates the probability that an individual is in the same state (either asleep or awake) at any given clock time in consecutive days across the study recording (Fig. 3a, Table S3). We compared the SRI between housed and unhoused participants in each season through a one-way ANOVA followed by Dunnett’s test. During the summer, community had a significant effect (p = 0.0456, Table S3.a); however, Dunnett’s tests revealed no differences in SRI between the housed and any of the unhoused groups (Table S3.b). Similarly, during the winter, there was a significant effect of community (p < 0.0001, Table S3.a), with housed participants displaying higher SRI (75 ± 2) than participants in the continuous-occupancy shelter (63 ± 4), tent city (56 ± 5), and tiny houses (61 ± 3) (Table S3.b). The overnight shelter (73 ± 2) had a similar SRI to the housed community, most likely attributable to the strict schedule of the shelter.

**Figure 3.**
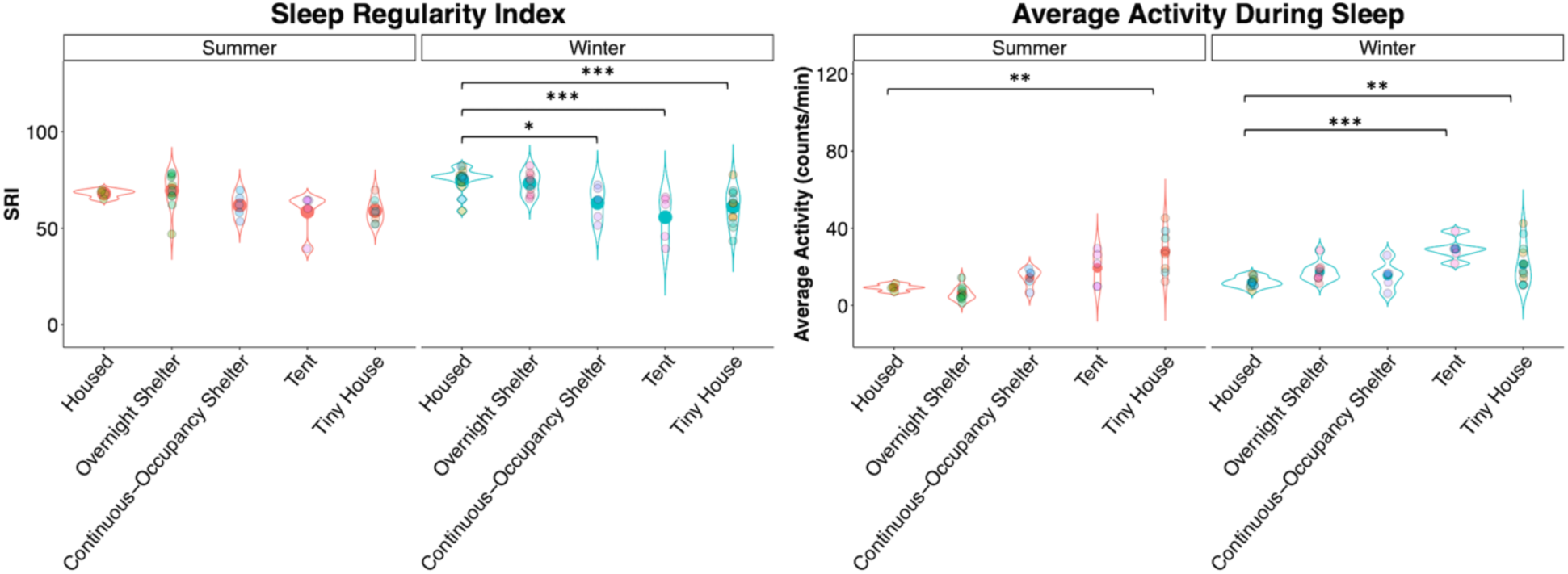
Violin plots of sleep regularity index and locomotor activity during sleep bouts within housed and unhoused communities during summer and winter. Large points in the middle of the plots represent the overall average SRI and average activity values. Smaller points represent individual participant values. Asterisks represent the results of Dunnett’s test: **p < 0.01, ***p < 0.001. See Tables S3 and S4 for ANOVA and Dunnett’s test results.

Second, we measured the average locomotor activity during the daily sleep bout (Fig 3b, Table S4) as an indicator of restless, low sleep quality. One-way ANOVA for the summer showed an effect of community (p < 0.0001, Table S4.a), with housed participants having less activity (10 ± 1 counts/min) than tiny house participants (28 ± 4; Dunnett’s test, Table S4.b). During the winter, there was also an effect of community type (p = 0.0003, Table S4.b), with housed participants (12 ± 1) displaying less activity during sleep than tent (29 ± 3) and tiny house (21 ± 3) participants (Table S4.b).

Lastly, because high levels of day-to-day variability in sleep timing indicate lower sleep quality, we compared the intraindividual variability of sleep parameters (represented by the standard deviation of each sleep parameter measured throughout days for each participant) across days for each community (Fig. 4, Table S5). During the summer, the variability in sleep parameters did not show a significant effect of community. During the winter, variability in sleep onset (p = 0.0040), midsleep (p = 0.0256), and duration (p = 0.0050) showed a significant effect of community (Table S5.a). Dunnett’s post-hoc comparisons revealed that the variability in sleep onset (p = 0.0051) and midsleep (p = 0.0332) was higher in tent participants than in housed participants (Table S5.b). Furthermore, sleep duration variability was higher for participants in the continuous-occupancy shelters (p = 0.0495), tents (p = 0.0113), and tiny houses (p = 0.0109) than for housed participants (Table S5.b).

**Figure 4.**
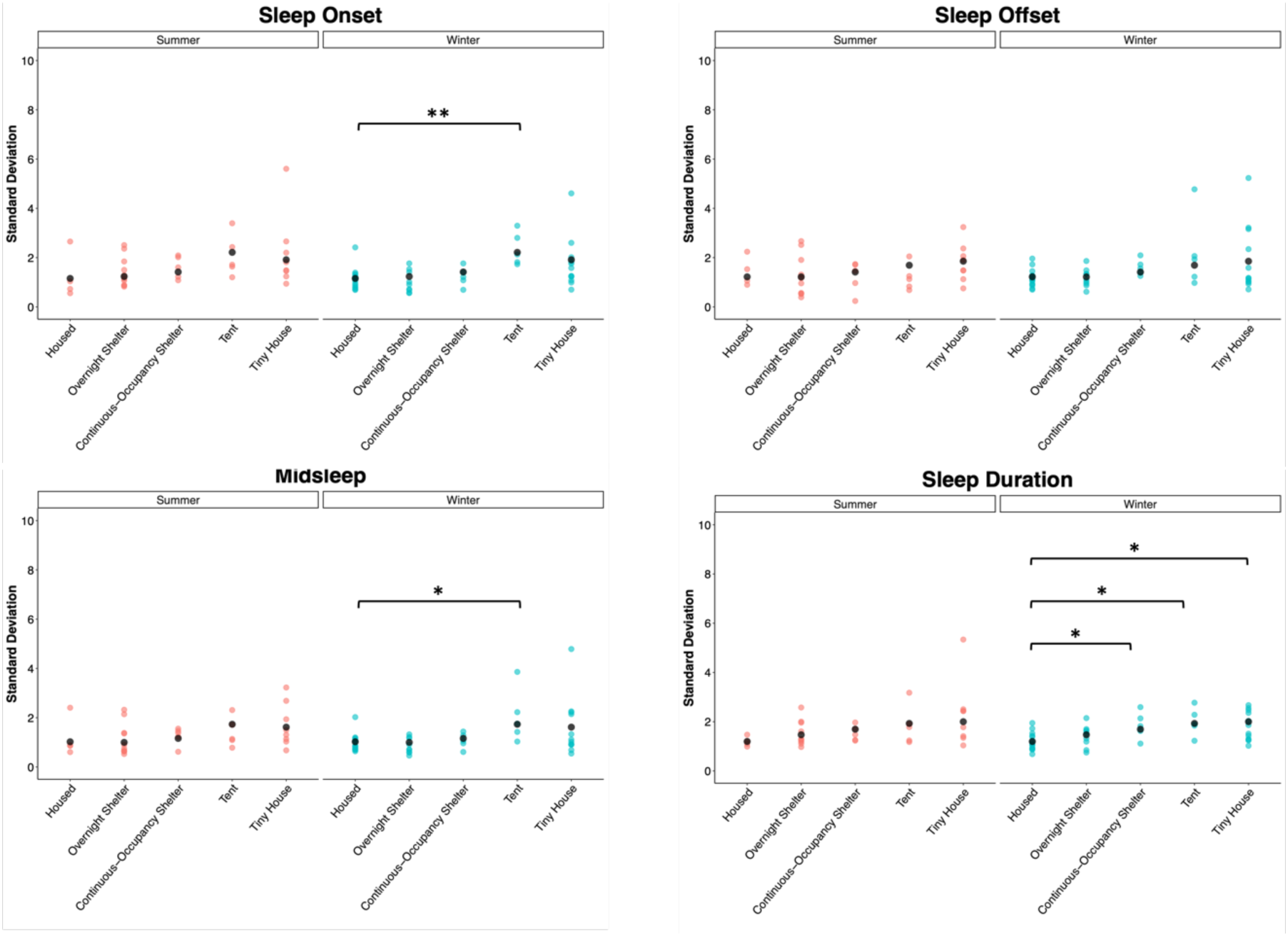
Intraindividual variation of sleep onset, offset, midsleep, and duration. Each point represents each participant’s standard deviation (all days combined), and black dots represent the mean standard deviation for each group. Asterisks represent the results of Dunnett’s test: *p < 0.05, **p < 0.01, ***p < 0.001. See Table S5 for ANOVA and Dunnett’s test results.

### Light Exposure

Light exposure is a critical modulator of sleep and is associated with sleep timing, and its seasonal changes are associated with sleep timing changes in people living in Seattle^11^. To determine the timing of light exposure during the day, we measured each participant’s first (morning) and last (evening) time of exposure to light above 50 lux (Fig. 5, Table S6). The 50-lux threshold was chosen as a likely threshold for circadian responses. A linear model with mixed effects for the first 50-lux light exposure showed an effect of season (p = 0.0044; Summer = 08:11 ± 00:13, Winter = 09:04 ± 00:16), community (p = 0.0001), and gender (p = 0.0052; Female = 08:10 ± 00:14, Male = 08:48 ± 00:15, NB/Trans/Other = 09:38 ± 00:42). Tent city participants were first exposed to 50-lux light 2h 54min earlier in the summer versus the winter (08:24 ± 00:17 during summer, 11:18 ± 00:44 during winter, p = 0.0328). The last 50-lux exposure had an effect of season (p = 0.0001; Summer = 10:08 ± 00:22, Winter = 08:24 ± 00:26) and community (p < 0.0001). The daily duration of exposure to at least 50-lux light showed a significant effect of season (p < 0.0001; Summer = 14h 04min ± 00:27, Winter = 11h 24min ± 00:38), community (p <0.0001), and gender (p = 0.0188; Female = 12h 48min ± 00:39, Male = 12h 32min ± 00:35, NB/Trans/Other = 11h 9min ± 01:53). The tent community was exposed daily to 50-lux light 6h33min more during the summer than the winter (10h 43min ± 00:16 during summer, 4h 10min ± 01:09 during winter; p = 0.0076). These results reveal that participants in the tent city were more exposed to natural daylight changes than those in other communities.

**Figure 5.**
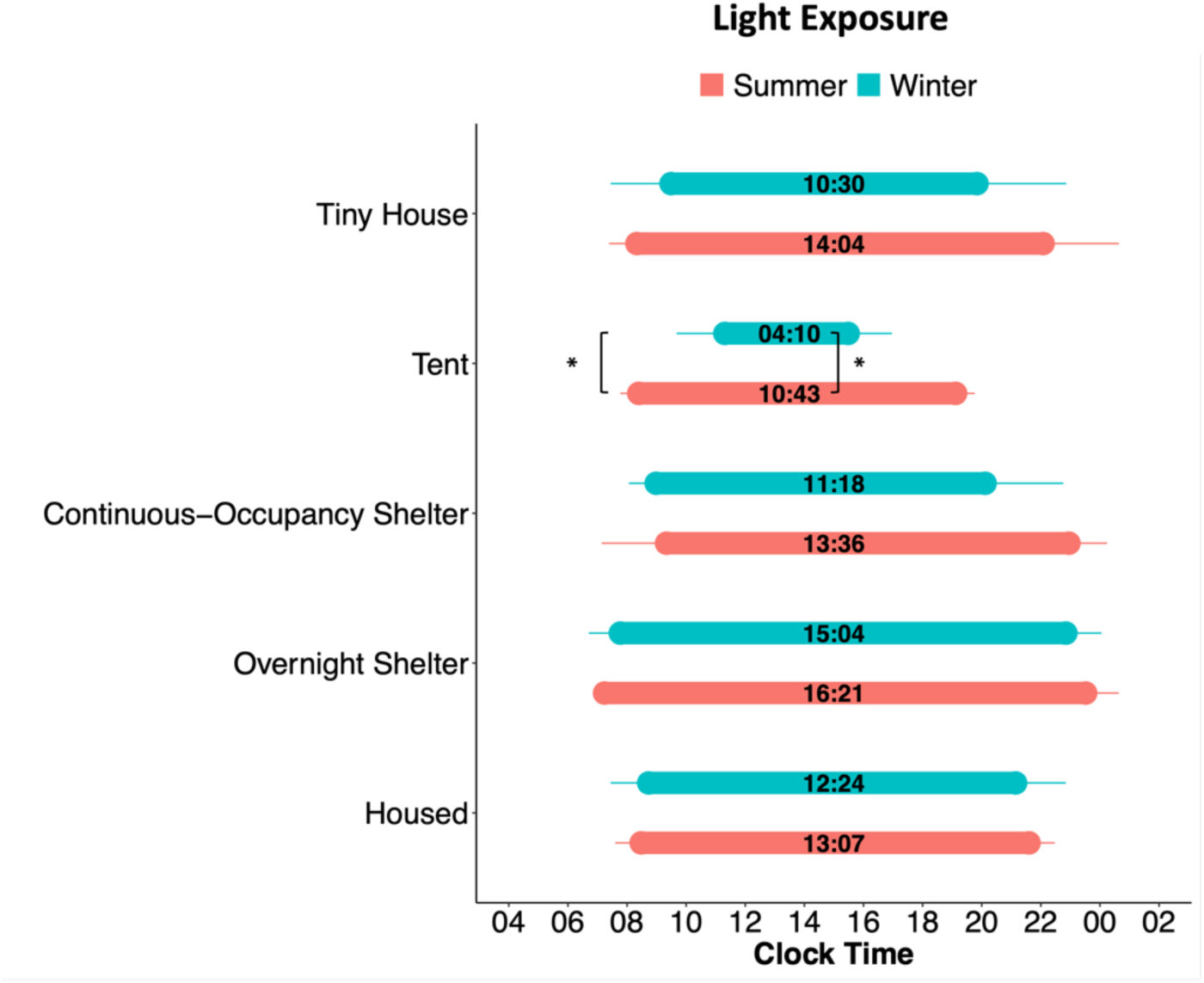
Timing of light exposure between communities and seasons. Bar plots of the average time of first exposure to 50 lux, last exposure to 50 lux, and light exposure duration (number within bars). Light exposure plots for each community are split between seasons. Error bars represent standard deviation. Symbols represent the results of a mixed-effects linear model followed by post hoc Tukey comparisons between summer and winter: *p < 0.05.

## Discussion

We show that actimetry-based sleep represents a sensitive method to detect differences in the timing and quality of sleep in four types of homeless shelter environments. For instance, the different homelessness environments are not associated with significant differences in sleep duration during the summer but are associated with substantial sleep disparities during the winter, when unhoused participants slept more than an hour and a half less per day than housed participants. Recording actimetry-based sleep represents a non-invasive, objective quantitative method that can reliably assess the effectiveness of interventions designed to improve the sleep of unhoused individuals. Given the importance of sleep to health and well-being, our sleep measurements should also serve as an indirect assessment of the health benefits of specific interventions.

Although the goal for any community should be that all people experiencing homelessness find stable housing, it is also essential to assess the effectiveness of transitional interventions.

Individually recorded, objective sleep data are extremely powerful as a metric for overall well-being, but it is critical to place results for each sleep parameter in each environmental context. The recommended sleep duration for the ages in our study (18-65 years, Table S7) is 7-9 hours^12^, and housed participants clearly met this recommendation. In our study, participants in overnight and continuous-occupancy shelters, and tiny houses met the 7-hour minimum during the winter.

However, they all slept much less than housed individuals, although the effect size was smaller in tiny houses. Unexpectedly, tent city participants did not sleep less than housed individuals in the winter, and this leads to the somehow counterintuitive impression that sleeping in a tent in the Seattle winter is similar to sleeping in a house. On the other hand, tent city participants displayed higher intra-individual variability in their sleep parameters and lower SRI, pointing to a lower day-to-day sleep consistency. Thus, an apparently sufficient sleep duration in tent city participants may not necessarily reveal adequate sleep, particularly considering the association between irregular sleep timing and adverse health outcomes^13–17^.

Our intraindividual and SRI analyses yielded no differences between housed individuals and participants in the overnight shelter. Although this suggests sleep in overnight shelters could be as good as in stable housing, this could instead be a result from rigid entry and exit schedules imposed on overnight shelter users.

Actimetry-based sleep has been used as a gold standard for field sleep studies, in which polysomnography is not feasible —particularly in homeless communities. Nevertheless, the main limitation of actimetry-based sleep is that sleep parameters come from a proxy of sleep: the lack of wrist activity. Overnight shelters have a very regimented schedule for arrival and bedtime as well as for wakeup time and departure. This is demonstrated by our waveform analysis, which reveals evening and morning locomotion peaks in overnight shelter participants. Thus, the apparent sleep hygiene in these participants recorded may simply reveal enforced inactivity.

As expected, light exposure measures revealed seasonal differences in tent-dwelling participants, but not in other communities. At 47° latitude, Seattle has substantial photoperiod differences between winter and summer, producing a nearly 8-hour sunrise-sunset time difference between summer and winter. Additionally, the sky is typically gloomy and overcast during the winter, even in daylight. In another study, the number of daylight hours in Seattle was the main predictor for sleep phase in housed young adults^11^, and was associated with a delayed sleep phase during the winter. Tent city participants experienced the seasonal difference in light exposure, but not in sleep timing, suggesting their environment prevents the expression of the effects of light on sleep phase.

Some limitations in our study should be noted. First, while the winter recordings reveal a highly negative influence of homelessness on sleep, we cannot assess which major environmental predictors of poor sleep are responsible —e.g., photoperiod, ambient temperature, or precipitation. Second, even though our housed controls were recorded during the same seasons as our unhoused subjects, the recordings were not done in the same year. Nevertheless, the differences between housed and unhoused participants clearly aligned with the pre-specified hypothesis that housed participants would sleep better. Third, our study does not present any health outcomes that could be accounted for by the differences in sleep between communities. Longitudinal studies could assess the predictive value of objectively recorded sleep parameters for health outcomes in unhoused people. Finally, our sample sizes for each shelter condition are relatively small, and our statistical analysis does not have the power to account for differences in participant age, time spent in each shelter condition, individual circumstances leading to being unhoused, substance usage, medications, or chronic health disorders that could influence sleep quality regardless of the shelter condition. Nevertheless, the effects we report demonstrate that the objective measurement of sleep represents a good instrument to assess the impact of specific interventions aimed at improving the well-being of people experiencing homelessness. Sleep and health have a bidirectional relationship in which poor sleep worsens health, and worse health, in turn, leads to poorer sleep. Accordingly, sleep disparities have lately received attention as a critical factor that perpetuates health inequities^18–20^, and evidence for this relationship has been documented through surveys of people experiencing homelessness^4,7^. Our study confirms that the current homelessness situation represents a human societal crisis that, beyond exposing people to the inhumane reality of not having stable housing, exposes them to a tremendous sleep and health disparity.

Sleep disparities among homeless populations were part of the arguments in the *Grants Pass vs. Johnson* case, in which the US Supreme Court ruled that unhoused people could be punished with fines or jail time for sleeping in public areas. In her dissenting opinion, Justice Sotomayor argued that “Sleep is a biological necessity, not a crime.”, an opinion supported by a vast body of scientific literature. Our study shows that people experiencing homelessness experience shorter and worse sleep; the stress of knowing that one can be arrested for sleeping in public areas will likely worsen these sleep parameters.

## Methods

### Participants

In partnership with organizations in the Seattle area that offer shelter to houseless people, we found that participants were eager to participate in our study and were compliant with requests to wear the actimetry watches and record their experiences in the diaries. We recruited participants from four types of unhoused communities to the study and, a convenience sample of housed University of Washington (UW) biology department students, staff and faculty. Detailed participant information is included in Table S7.

The four Seattle-based homeless advocacy and assistance organizations we worked with were known to our network and agreed to participate through trusted relationships. ROOTS is a youth shelter in the University District that offers emergency overnight shelter for young adults aged 18-25. A waitlist is assembled each evening between 20:00 and 20:30. ROOTS’ website explains that “if more than 45 people are trying to access the shelter, a random drawing is done at 8:30 pm to determine who will be turned away.” At the time of the study, volunteers were on site to make and serve breakfast each morning, and people were expected to leave by 08:30 and return the following evening if they wanted to stay another night. We included 9 ROOTS participants in the winter (2019) and 10 in the summer (2018).

Nickelsville organizes two tiny house communities with city permission, one in the University District (with 19 tiny homes) and one east of downtown in the “Central District” of Seattle (with 14 homes). These self-governed, democratically organized villages are established on spaces about the size of a city lot, each with a couple of dozen shed-sized wooden structures (2.4x3.6 m). Each tiny home has a window and includes a bed, light and a heater. On-site shared toilets, shower rooms, and laundry machines are available. People may leave their belongings in their locked units, may have pets, and can stay with a partner and/or child. Dinners are eaten communally in an open kitchen. Nickelsville chose to participate in our study after the residents in each location voted. Between the two sites, we included 14 in the winter of 2023 and 10 in the summer of 2022.

SHARE has organized self-governed, democratically organized tent cities (with up to 100 participants each) around Seattle since 1990. Tents are set up in rows on the site, and individuals, couples, families, and pets are welcome. Typically, each tent city must move to a new location every 90 days; when our study was conducted, Tent City 3 was established on the St. Mark’s Cathedral parking lot (Summer 2022) and relocated to a University of Washington parking lot the following winter. Tents are set up on a wooden pallet, and several portable toilets are on site; there is also a minimally equipped kitchen area where communal meals can be taken. Tent cities are sanctioned, so they are not at risk of being disturbed by police. Like Nickelsville, these tent cities welcome participants who agree to the inclusion terms (e.g., contribute hours of service, agree not to bring drugs or alcohol on site, and attend weekly encampment meetings). Like Nickelsville, participation in our study was agreed to after a vote of residents at their weekly meeting. We included 9 subjects in the winter of 2023 and 12 in the summer of 2022.

Among many other efforts, SHARE also operates a 24-hour, self-managed, semi-private (shared room) shelter called the “Bunkhouse.” There is no maximum length of stay at the Bunkhouse.

Seven Bunkhouse participants were in the study in the summer of 2022, and 7 in the winter of 2023.

All participants signed informed consent forms and each participant who completed their daily sleep diaries and wore wrist actimeters as requested was compensated with weekly monetary gift cards. All studies were approved by the UW Human Subjects Division.

### Data Collection

All activity and light exposure data was recorded using Actiwatch Spectrum Plus loggers (Phillips, Respironics, Bend, OR) programmed to collect data in 1-min epochs (except for summer overnight shelter recordings that collected data in 15-sec epochs). Participants also completed a demographic form (in person) and daily sleep logs using online forms asking their bedtime, wake time, nap time (if taken), and where they slept the previous night. In the summer, we recorded participants for 5 weeks except for the overnight shelter participants, who were recorded for 4 weeks, and housed participants, who were recorded for 2 weeks. In the winter, recordings lasted 4 weeks except in housed individuals, lasting 7 weeks.

We collected data from two Seattle tiny house locations, but data were analysed together since a t-test showed no difference between sleep parameters.

All preparation, analysis, and data plotting were performed using R studio version 2023.06.1 + 524 unless otherwise indicated. A p-value of <0.05 was considered statistically significant.

### Inclusion Criteria

Individual actograms were inspected with Philips Actiware (version 6.09) software to check for watch malfunction and to see that watches were worn for an appropriate number of days during recording. Individual sleep diaries were also inspected and compared to Actiwatch recordings. Any dates with more than a 3-hour discrepancy in watch-determined sleep or wake timing compared to self-reported sleep dairy sleep or wake timing were excluded. Any night on which a sleep event was not detected from actigraphy due to locomotor activity and light exposure measurements, and for which no sleep diary entry was provided, was considered a sleepless night.

Dates on which subjects did not sleep at their typical site were excluded. For example, if a tent city participant slept at a hotel on one particular night, that date was removed from the analysis. Participants with less than 14 nights of sleep recordings were omitted. Also, participants with overnight shift jobs were excluded. The detail of the 22 excluded participants is included in Table S7.

### Sleep Parameters

Actiwatch recordings were downloaded and exported using the Philips Actiware software. Using activity measurements, the software estimates sleep parameters including onset, offset, and duration.

#### Statistical analysis

Each participant’s daily sleep parameters (sleep onset, offset, duration, and midsleep timing) were averaged for all days, generating one value per participant. Data were analysed using a linear model for each season (winter or summer) and sleep parameter separately, considering gender (female, male, non-binary, and transgender) and sleeping community as factors. We used linear model coefficient values to compare housed sleep parameter values to all other unhoused (tent, tiny home, continuous-occupancy shelter, overnight shelter) values.

To assess differences in intraindividual variance of sleep parameters between communities, we calculated the standard deviation of each sleep parameter across all days recorded for each participant. Then, we used a separate one-way ANOVA per season to compare intraindividual variances between communities. Lastly, a Dunnett’s test using the Desctools package was run to compare intraindividual variances of each unhoused community to housed community variance^21^.

### Light Exposure

All light exposure data were analysed using the Actiwatch white-light lux reading. We determined the first, the last, and the duration of daily exposure times to a specific light intensity using the 1-minute epoch raw light data for each individual. We analysed the mean time of the first and last exposure to a 50-lux light intensity for each participant. Importantly, 50 lux is not necessarily a circadian threshold for non-visual responses^22,23^, but instead it was chosen as an arbitrary value that would likely be above the threshold for circadian photic stimulation for most of the participants as previously described^11,24,25^.

#### Statistical analysis

Differences in light exposure times between participants in all communities and seasons were analysed using a linear mixed effects model (LMEM) using the lme4 package^26^. QQ plots for every LMEM fit were first analysed to check normality of the residues of the model. The model was run on each light timing variable (first 50-lux time, last 50-lux time, and total duration under 50-lux) separately considering environment (housed, continuous-occupancy shelter, overnight shelter, tiny house and tents), season (summer or winter), and gender as factors. Tukey post hoc comparisons were used to determine differences in the timing of light exposure between communities and seasons.

### Waveforms

To determine differences in overall activity, raw 1-minute activity data for each participant was binned into means of 30-min intervals. After 30-min data were log-transformed, activity was smoothed by a one-hour running average. The resulting data generated individual 24-hour activity waveforms for each participant. Individual waveforms were in turn used to generate mean waveforms, leading to two activity waveforms per community (one for each season).

#### Statistical analysis

Analysis of summer and winter waveforms between sleeping communities was done using a two-way mixed ANOVA with community as a between-participants factor and time as a within-participants factor. We compared community activity during specific time windows during the morning (summer = 7:00-10:00; winter = 7:00-9:00) or evening (summer = 19:00-21:30; winter = 20:00 – 22:00). For these time window analyses, we added cumulative activity counts for each participant and used a one-way ANOVA to compare values between communities. Time windows for morning and evening analysis were chosen based on visual inspection and the slope of the waveforms.

### Sleep Regularity Index

To determine sleep regularity of participants, we calculated their sleep regularity index (SRI), as a measure of the probability of an individual being in the same state (asleep vs. awake) at any two time points 24 hours apart^27^. An individual who falls asleep and wakes up at exactly the same time every day will score a 100; an individual who sleeps and wakes randomly from day to day will score a 0.

#### Statistical analysis

SRI values were calculated using daily sleep onset and offset timing determined by the Actiwatch. SRI data were analysed using a separate one-way ANOVA and Dunnett’s test per season, similar to the analysis done with the intraindividual variance.

### Activity During Sleep

Average activity during sleep was determined using the Actiwatch raw activity data and daily sleep onset and offset data for each participant. Activity counts/min during the watch-determined sleep bouts were averaged per day for each subject, and daily 0mean values were averaged for each subject, leading to one mean value for each participant. Mean values were then calculated per community separated by seasons.

#### Statistical analysis

We used a separate one-way ANOVA and Dunnett’s test per season, similar to the analysis done with the intraindividual variance.

## Data availability

All data used to generate figures will be deposited in GitHub with the subdomain to be named upon acceptance of the manuscript.

## Acknowledgments

We thank the homeless advocacy organizations that agreed to work with us: SHARE, Nickelsville, ROOTS, along with all the individual participants who volunteered their time and personal sleep data. We are particularly grateful to Michele Marchand (SHARE/WHEEL Organizer) and Kat Ousley (ROOTS Shelter Director). We also thank Hannah Hinton and Olivia Sather for their assistance with data collection. This Study was supported by NIH R01HL162311, a grant from the University of Washington Population Health Initiative, and the Department of Biology and the Center for Studies in Demography and Ecology at University of Washington. ZWA was also partially supported on NSF CAREER grant (SES-2142964).

## Supplementary information

**Table S1.a.**
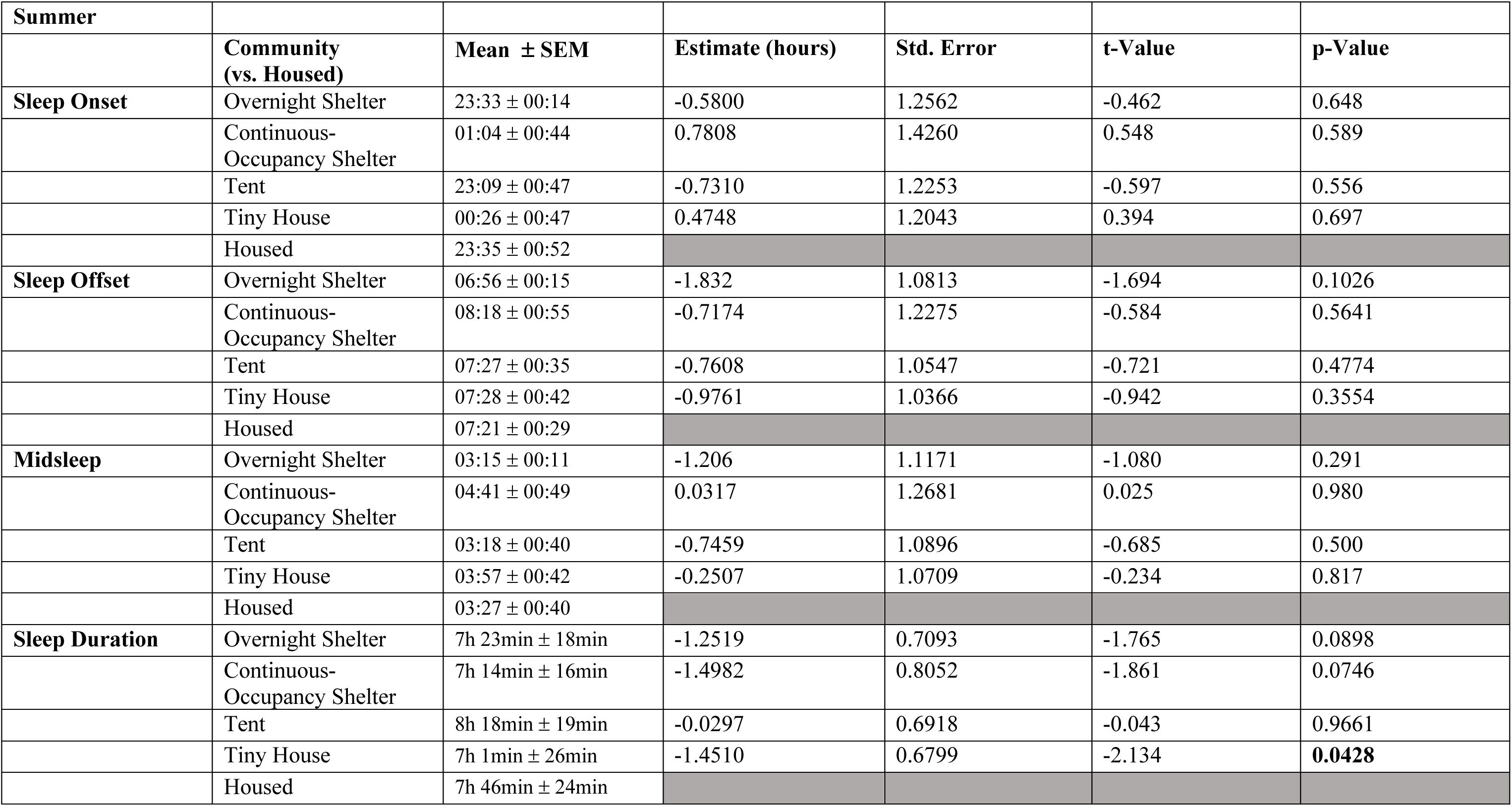
Sleep parameters linear model estimates for summer.

| Summer |  |  |  |  |  |  |
| --- | --- | --- | --- | --- | --- | --- |
|  | **Community**<br>**(vs. Housed)** | **Mean ± SEM** | **Estimate (hours)** | **Std. Error** | **t-Value** | **p-Value** |
| **Sleep Onset** | Overnight Shelter | 23:33 ± 00:14 | -0.5800 | 1.2562 | -0.462 | 0.648 |
|  | Continuous-Occupancy Shelter | 01:04 ± 00:44 | 0.7808 | 1.4260 | 0.548 | 0.589 |
|  | Tent | 23:09 ± 00:47 | -0.7310 | 1.2253 | -0.597 | 0.556 |
|  | Tiny House | 00:26 ± 00:47 | 0.4748 | 1.2043 | 0.394 | 0.697 |
|  | Housed | 23:35 ± 00:52 |  |  |  |  |
| **Sleep Offset** | Overnight Shelter | 06:56 ± 00:15 | -1.832 | 1.0813 | -1.694 | 0.1026 |
|  | Continuous-Occupancy Shelter | 08:18 ± 00:55 | -0.7174 | 1.2275 | -0.584 | 0.5641 |
|  | Tent | 07:27 ± 00:35 | -0.7608 | 1.0547 | -0.721 | 0.4774 |
|  | Tiny House | 07:28 ± 00:42 | -0.9761 | 1.0366 | -0.942 | 0.3554 |
|  | Housed | 07:21 ± 00:29 |  |  |  |  |
| **Midsleep** | Overnight Shelter | 03:15 ± 00:11 | -1.206 | 1.1171 | -1.080 | 0.291 |
|  | Continuous-Occupancy Shelter | 04:41 ± 00:49 | 0.0317 | 1.2681 | 0.025 | 0.980 |
|  | Tent | 03:18 ± 00:40 | -0.7459 | 1.0896 | -0.685 | 0.500 |
|  | Tiny House | 03:57 ± 00:42 | -0.2507 | 1.0709 | -0.234 | 0.817 |
|  | Housed | 03:27 ± 00:40 |  |  |  |  |
| **Sleep Duration** | Overnight Shelter | 7h 23min ± 18min | -1.2519 | 0.7093 | -1.765 | 0.0898 |
|  | Continuous-Occupancy Shelter | 7h 14min ± 16min | -1.4982 | 0.8052 | -1.861 | 0.0746 |
|  | Tent | 8h 18min ± 19min | -0.0297 | 0.6918 | -0.043 | 0.9661 |
|  | Tiny House | 7h 1min ± 26min | -1.4510 | 0.6799 | -2.134 | **0.0428** |
|  | Housed | 7h 46min ± 24min |  |  |  |  |

**Table S1.b.** Sleep parameters linear model estimates for winter.

| Winter |  |  |  |  |  |  |
| --- | --- | --- | --- | --- | --- | --- |
|  | **Community**<br>**(vs. Housed)** | **Mean ± SEM** | **Estimate (hours)** | **Std. Error** | **t-Value** | **p-Value** |
| **Sleep Onset** | Overnight Shelter | 23:28 ± 00:17 | 0.2194 | 0.6396 | 0.343 | 0.7335 |
|  | Continuous-Occupancy Shelter | 23:50 ± 00:25 | 0.3264 | 0.8499 | 0.384 | 0.7032 |
|  | Tent | 23:11 ± 1:06 | -0.1691 | 0.7639 | -0.221 | 0.8260 |
|  | Tiny House | 23:54 ± 00:33 | 0.7926 | 0.5815 | 1.363 | 0.1814 |
|  | Housed | 23:07 ± 00:19 |  |  |  |  |
| **Sleep Offset** | Overnight Shelter | 06:28 ± 00:20 | -1.2654 | 0.6851 | -1.847 | 0.073 |
|  | Continuous-Occupancy Shelter | 06:41 ± 00:27 | -0.8939 | 0.9104 | -0.982 | 0.333 |
|  | Tent | 06:59 ± 1:12 | -0.6638 | 0.8183 | -0.811 | 0.423 |
|  | Tiny House | 07:03 ± 00:36 | -0.5572 | 0.6230 | -0.894 | 0.377 |
|  | Housed | 07:31 ± 00:12 |  |  |  |  |
| **Midsleep** | Overnight Shelter | 02:58 ± 00:17 | -0.5230 | 0.6369 | -0.821 | 0.4170 |
|  | Continuous-Occupancy Shelter | 03:15 ± 00:24 | -0.2837 | 0.8464 | -0.335 | 0.7394 |
|  | Tent | 03:06 ± 1:09 | -0.4164 | 0.7607 | -0.547 | 0.5875 |
|  | Tiny House | 03:29 ± 00:34 | 0.1177 | 0.5791 | 0.203 | 0.8401 |
|  | Housed | 03:19 ± 00:14 |  |  |  |  |
| **Sleep Duration** | Overnight Shelter | 6h 59min ± 15min | -1.4848 | 0.3663 | -4.053 | **0.0003** |
|  | Continuous-Occupancy Shelter | 6h 50min ± 23min | -1.2202 | 0.4868 | -2.507 | **0.0168** |
|  | Tent | 7h 48min ± 7min | -0.4948 | 0.4375 | -1.131 | 0.2656 |
|  | Tiny House | 7h 9min ± 15min | -1.3498 | 0.3331 | -4.052 | **0.0003** |
|  | Housed | 8h 24min ± 16min |  |  |  |  |

**Table S2.** Waveform morning and evening cumulative counts one-way ANOVA.

| **Summer** |  |  |  |  |
| --- | --- | --- | --- | --- |
|  | **Fixed Effect** | **Df** | **F-value** | **p-value** |
| Morning<br>(7:00-10:00) | Community | 4 | 2.9 | **0.0406** |
| Evening<br>(19:00-21:30) | Community | 4 | 2.584 | 0.0596 |

| **Winter** |  |  |  |  |
| --- | --- | --- | --- | --- |
|  | **Fixed Effect** | **Df** | **F-value** | **p-value** |
| Morning<br>(7:00-9:00) | Community | 4 | 4.365 | **0.0051** |
| Evening<br>(20:00-22:00) | Community | 4 | 2.068 | 0.1030 |

**Table S3.a.** SRI one-way ANOVA.

|  | **Fixed Effect** | **Df** | **F-value** | **p-value** |
| --- | --- | --- | --- | --- |
| **Summer** | Community | 4 | 2.804 | **0.0456** |
| **Winter** | Community | 4 | 8.579 | **< 0.0001** |

**Table S3.b.** Dunnett’s Test for SRI.

|  | **Community (vs. Housed)** | **p-value** |
| --- | --- | --- |
| **Summer** | Overnight Shelter | 0.9896 |
|  | Continuous-Occupancy Shelter | 0.5819 |
|  | Tent | 0.2461 |
|  | Tiny House | 0.2139 |
| **Winter** | Overnight Shelter | 0.9705 |
|  | Continuous-Occupancy Shelter | **0.0327** |
|  | Tent | **0.0003** |
|  | Tiny House | **0.0005** |

**Table S4.a.** Activity During Sleep one-way ANOVA.

|  | **Fixed Effect** | **Df** | **F-value** | **p-value** |
| --- | --- | --- | --- | --- |
| **Summer** | Community | 4 | 10.7 | **< 0.0001** |
| **Winter** | Community | 4 | 6.791 | **0.0003** |

**Table S4.b.** Dunnett’s Test for Activity During Sleep.

|  | **Community (vs. Housed)** | **p-value** |
| --- | --- | --- |
| **Summer** | Overnight Shelter | 0.8665 |
|  | Continuous-Occupancy Shelter | 0.6534 |
|  | Tent | 0.1342 |
|  | Tiny House | **0.0011** |
| **Winter** | Overnight Shelter | 0.2550 |
|  | Continuous-Occupancy Shelter | 0.7850 |
|  | Tent | **< 0.0001** |
|  | Tiny House | **0.0063** |

**Table S5.a.** Intraindividual variance one-way ANOVA.

| **Summer** |  |  |  |  |
| --- | --- | --- | --- | --- |
|  | **Fixed Effect** | **Df** | **F-value** | **p-value** |
| Sleep Onset | Community | 4 | 1.09 | 0.3813 |
| Sleep Offset | Community | 4 | 0.882 | 0.4877 |
| Midsleep | Community | 4 | 1.599 | 0.5395 |
| Sleep Duration | Community | 4 | 1.379 | 0.2033 |

| **Winter** |  |  |  |  |
| --- | --- | --- | --- | --- |
|  | **Fixed Effect** | **Df** | **F-value** | **p-value** |
| Sleep Onset | Community | 4 | 4.544 | **0.0040** |
| Sleep Offset | Community | 4 | 2.09 | 0.1001 |
| Midsleep | Community | 4 | 3.109 | **0.0256** |
| Sleep Duration | Community | 4 | 1.0514 | **0.0049** |

**Table S5.b.** Dunnett’s test for intraindividual variance analysis.

| **Winter** |  |  |
| --- | --- | --- |
|  | **Community (vs. Housed)** | **p-value** |
| **Sleep Onset** | Overnight Shelter | 0.9892 |
|  | Continuous-Occupancy Shelter | 0.9965 |
|  | Tent | **0.0051** |
|  | Tiny House | 0.0988 |
| **Sleep Offset** | Overnight Shelter | 1.0000 |
|  | Continuous-Occupancy Shelter | 0.7848 |
|  | Tent | 0.1168 |
|  | Tiny House | 0.1583 |
| **Midsleep** | Overnight Shelter | 0.9898 |
|  | Continuous-Occupancy Shelter | 0.9982 |
|  | Tent | **0.0332** |
|  | Tiny House | 0.1764 |
| **Sleep Duration** | Overnight Shelter | 0.7306 |
|  | Continuous-Occupancy Shelter | **0.0495** |
|  | Tent | **0.0113** |
|  | Tiny House | **0.0109** |

**Table S6.** 50-lux exposure times linear model estimates.

|  | **Fixed Effect** | **Df** | **F-value** | **p-value** |
| --- | --- | --- | --- | --- |
| First Exposure | Season | 1 | **8.7341** | **0.0044** |
|  | Community | 4 | 6.9813 | **0.0001** |
|  | Gender | 3 | 4.6709 | **0.0052** |
|  | Season:Community | 4 | 2.2880 | 0.0697 |
| Last Exposure | Season | 1 | 17.1610 | **0.0001** |
|  | Community | 4 | 15.9751 | **< 0.0001** |
|  | Gender | 3 | 1.6547 | 0.1858 |
|  | Season:Community | 4 | 1.6711 | 0.1678 |
| Duration | Season | 1 | 21.2113 | **< 0.0001** |
|  | Community | 4 | 17.3567 | **< 0.0001** |
|  | Gender | 4 | 3.5704 | **0.0188** |
|  | Season:Community | 4 | 2.3437 | 0.0643 |

**Table S7.** Participant characteristics.

|  | **Tiny House** |  | **Tent City** |  | **Overnight Shelter** |  | **Continuous-Occupancy Shelter** |  | **Housed** |  |
| --- | --- | --- | --- | --- | --- | --- | --- | --- | --- | --- |
|  | Winter<br>n = 14 | Summer<br>n = 10 | Winter<br>n = 9 | Summer<br>n = 12 | Winter<br>n = 9 | Summer<br>n = 10 | Winter<br>n = 7 | Summer<br>n = 7 | Winter<br>n = 14 | Summer<br>n = 6 |
| Recording dates | 1/23/2023-2/22/2023 | 9/5/2022-10/17/2022 | 1/23/2023-2/22/2023 | 9/5/2022-10/17/2022 | 2/7/2019-3/6/2019 | 9/5/2018-10/4/2018 | 1/23/2023-2/22/2023 | 9/5/2022-10/17/2022 | 2/15/2021-4/7/2021 | 9/26/2023-10/14/2023 |
| **Self-Reported Gender (%)** |  |  |  |  |  |  |  |  |  |  |
| Female | 36 | 30 | 33 | 33 | 22 | 10 | 0 | 14 | 43 | 67 |
| Male | 50 | 70 | 67 | 58 | 67 | 80 | 86 | 71 | 50 | 33 |
| Non-binary or transgender | 14 | 0 | 0 | 0 | 11 | 10 | 14 | 14 | 0 | 0 |
| Other or missing | 0 | 0 | 11 | 8 | 0 | 0 | 0 | 0 | 7 | 0 |
| **Self-Reported Race (%)\*** |  |  |  |  |  |  |  |  |  |  |
| Caucasian | 57 | 60 | 78 | 67 |  |  | 86 | 86 | 36 | 67 |
| Black | 35 | 20 | 0 | 56 |  |  | 14 | 14 | 14 | 0 |
| Asian | 0 | 10 | 0 | 0 |  |  | 0 | 0 | 7 | 17 |
| Hispanic | 0 | 0 | 0 | 0 |  |  | 0 | 0 | 35 | 17 |
| American Indian | 14 | 20 | 11 | 14 |  |  | 14 | 0 | 0 | 0 |
| Other or missing | 0 | 0 | 11 | 0 |  |  | 14 | 0 | 7 | 0 |
| **Age (years): mean [range]** | 43 [28-60] | 48 [31-63] | 44 [35-55] | 44 [24-65] | 22 [19-26] | 23 [18-26] | 43 [32-58] | 44 [26-58] | 38 [24-68] | 30 [22-58] |
| **Excluded participants (n)** | 2 | 2 | 4 | 7 | 1 | 0 | 2 | 2 | 0 | 2 |

\*Participants may have reported more than one race, so that the total percent could be greater than 100%
\*\* Overnight shelter race was not self-identified

**Figure S1.**
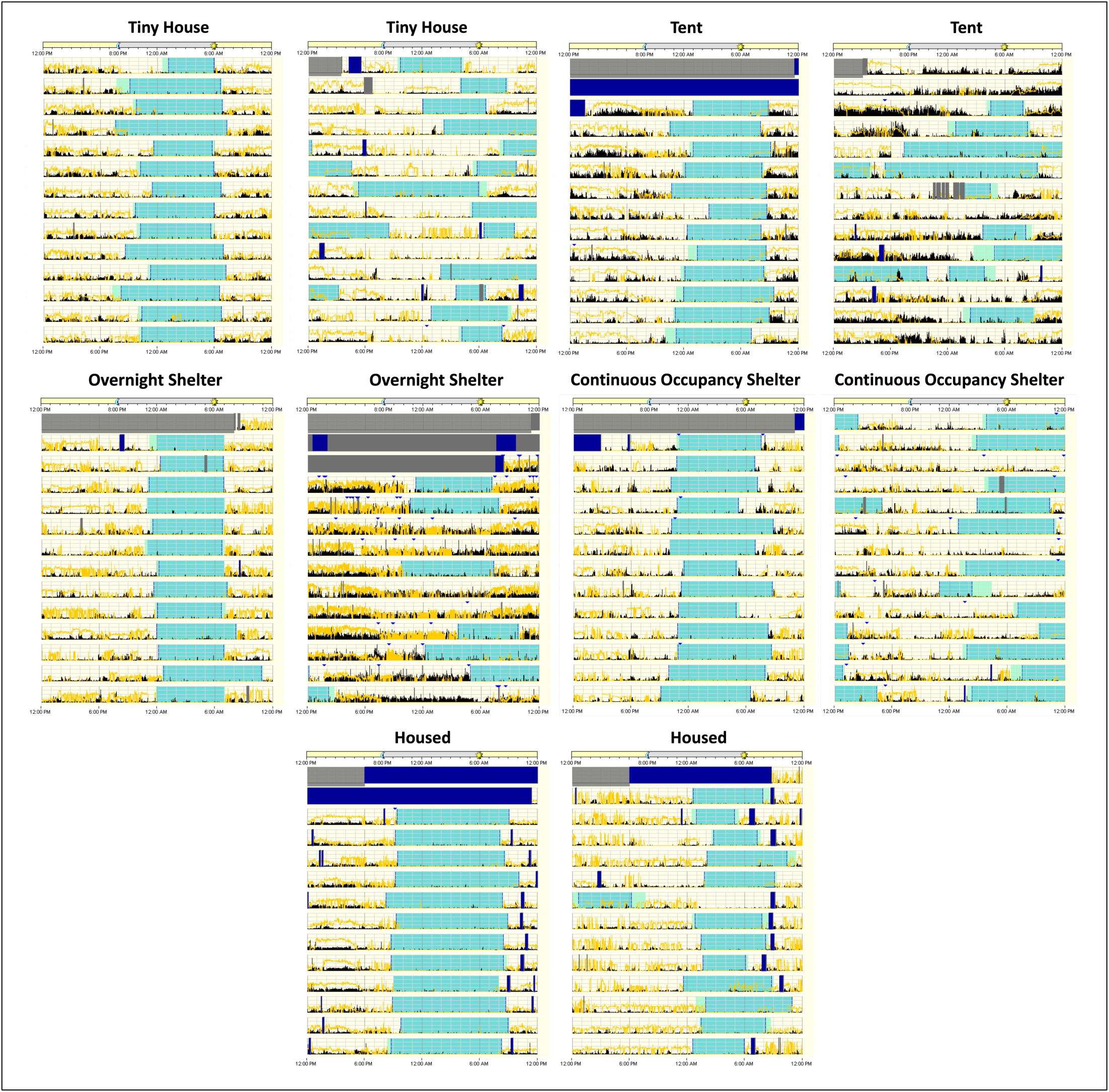
Actiwatch generated 24-hour actograms (12:00–12:00) of two participants from each community. The two actograms are representative examples from opposite ends of the spectrum between regular and most irregular sleep for each community. Sleep bouts are shown in light blue shaded sections, activity level is represented by black lines, and white light exposure by yellow lines.

